# Long-read single cell transcriptomics uncovers isoform preferences in developing human retina

**DOI:** 10.64898/2026.08.17.745278

**Authors:** L. Kaplan, J. Pang, T. A Reh

## Abstract

Retinal development has been extensively studied and key transcriptional regulators that drive fate decisions have been identified for major cell classes. These findings were confirmed and deepened in recent years with the advance of single cell RNA sequencing (Scrase). However, many processes that guide progenitor to postmitotic cell differentiation remain elusive, especially since some genes seem to yield different cell populations without apparent correlation with expression level or timing. Here, differential transcript isoform usage might play a role in diversifying the function of developmental genes. In short-read based scRNAseq, isoforms can only be identified if a read maps to a unique sequence or exon junction. However, due to the sparsity and very short reads, these events are extremely rare. We combined a commercial scRNAseq kit, that produces barcoded, full-length cDNA with Oxford Nanopore Technologies based long-read sequencing to generate the first single cell long-read sequencing dataset of fetal human retina. It can help elucidate the role of alternative splicing in retinal development and guide the design of transcript-specific gene therapies for retinal regeneration.

## Background and Summary

The embryonic development of the retina is a process conserved throughout vertebrates. All neuronal cell types and the Müller glia originate from a common progenitor and mature cells arise in well-defined order [1]. In the past decades considerable progress has been made to identify major genetic regulators that drive maturation and cell fate choice. More recent single cell transcriptome analyses of developing mouse and human retina have confirmed and extended the basic model of retinal histogenesis derived from experimental studies of model organisms. Transcriptomic data, complemented by single cell analyses of cis-regulatory regions (via ATACseq) of developing retinal cells has deepened our knowledge of the gene regulatory networks underlying cell fate decisions [2], [3], [4], [5].

However, one mechanism that is less explored is alternative splicing, in which the same gene locus produces a variety of mature mRNAs. Some outcomes influence mostly posttranscriptional regulation due to a change in untranslated regions (UTR) leading to altered mRNA stability, secondary structure, RNA binding protein attachment sites or availability of miRNA targeting sites. Additionally, many transcript isoforms differ in the number or length of translated exons producing truncated or conformationally changed mature proteins [6], [7].

Alternative splicing events have been studied during development of various tissues including the retina (comprehensively reviewed here [7], [8]) and for disease associated genes [9]. However, looking at the tissue level of the retina, especially during development, with its highly heterogenous cell composition and tight genetic regulation vastly underestimates cell type specific contributions. As noted above, single cell transcriptomics and epigenomics have provided a wealth of new data; however, due to their reliance on shallow, short-read sequencing, they are ill-suited to study the variation of transcript isoforms, as multiple splice site covering reads are necessary for proper identification. Long-read sequencing (LRS), pioneered by PacBio and Oxford Nanopore Technologies, have opened up new research opportunities and are now increasingly being used for this purpose [9], [10]. They can generate reads of hundreds of thousands of base pairs in length, covering not only multiple splice sites, but even complete transcripts. Existing datasets are focused either on bulk and/or adult tissue, but single cell developmental data is still lacking.

To fill this gap, we generated a single cell LRS based transcriptome dataset of fetal human retina that not only reproduces known typical cell type clustering and gene expression patterns but also reveals the fine-grained landscape of around 80000 transcript isoforms.

## Methods

### Isolation of retinal tissue

Human fetal eyes of gestational ages between 70 to 150 days were received from the Birth Defects Research Laboratory at the University of Washington following the approved protocol (UW5R24HD000836). After the removal of cornea and lens, retina was detached from RPE/choroid and separated into temporal and nasal sections. These were further dissected to retrieve central pieces around the presumptive fovea and peripheral parts distal to the optic nerve head on the nasal side.

### Generation of full-length cDNA libraries of single cells

For single-cell LRS we dissected temporal-central and nasal-peripheral retina of a gestational age 82 eye. Retinas were digested in Papain (Worthington) for 10 min at 37° C, followed by mechanical dissociation using a P1000 pipette. Ovomucin (Worthington) was added to the solution to stop the reaction. The suspension was passed through a 70 µm cell strainer and spun down for 7 min at 400 g, 4° C. Full length cDNA library was prepared using 10x Genomics’ Chromium Next GEM Single Cell 3’ Reagent Kit v3.1 (PN-1000268, 10x Genomics) according to manufacturer’s instructions up until the point of preamplification and before fragmentation.

Sequencing was performed at the Nanopore Sequencing Core of the University of Washington. Briefly, library size distribution was analyzed using a Femto Pulse (Agilent). Sequencing-ready samples were prepared using the Ligation Sequencing Kit V14 (SQK-LSK114, Oxford Nanopore) and PCR Expansion (EXP-PCA001, Oxford Nanopore) according to a published protocol [11]. The sequencing was run on a PromethION (Oxford Nanopore) and we received basecalled (dorado, Oxford Nanopore), unmapped bam files with ~100 million reads per sample (>20 000 reads per cell) and a mean read quality of >20. QC, alignment and counting of reads was performed using the epi2me-labs/wf-single-cell nextflow workflow (version 3.3.2 [12]) yielding cell by gene and cell by transcript count matrices. The used genome reference was GRCh38 with GENCODE v44/Ensembl 110 annotations retrieved from 10x Genomics (Human reference (GRCh38) - 2024-A, [13])

### Single cell pipeline with Seurat

Single cell transcriptomic analyses were performed using R programming [14] and the Seurat toolkit [15]. Cell by gene count matrices from each sample were initially processed separately. Briefly, we first filtered to include only high-quality cells/genes (min.cells = 50, min.features = 500, percent mitochondrial genes <10%). Resulting objects were then integrated using the RPCAIntegration method from which a common UMAP embedding was produced. Cell types were identified using known markers as previously described [16]. Filtered cells, genes, cell-labels and UMAP coordinates were transferred to the cell by transcript based object.

For coverage plots, we first split the reads of the aligned bam files (via samtools [17]) according to cell type and used them to generate strand specific bigWig files (deepTools bamCoverage [18]) representing pseudobulk coverage. BigWig files and the gene annotation (GTF) of the genome reference were used to plot coverage tracks across transcripts with the Gviz package [19].

### miRNA binding site identification

For global miRNA binding site predictions with TargetScan we first downloaded predicted target with context scores (Predicted_Targets_Context_Scores.default_predictions.txt, [20], [21]) and filtered to contain only conserved miRNAs and target transcript IDs detected in our single cell data.

Prediction with scanMIR [22] was available through an R package. It uses biochemical models for each miRNA (KdModels) to calculate miRNA-transcript interaction. We used KdModels for the previously defined conserved miRNAs as query and 3’UTRs from detected isoforms extracted from the GTF file of our reference as the targets. Only hits with a 6mer and 7mer binding and strong predicted repression (top 10 %) were kept.

As running PRIMITI [23], [24] was resource intensive for all detected isoforms, we limited the analysis to a subset of protein coding transcripts. Results were filtered to only include miRNA-transcript pairs with predicted interaction.

### PCR/qPCR

For RT-qPCR analysis retina was dissected in temporal-central and nasal-peripheral samples and RPE was peeled from the choroid and stored at −80 °C until further use. RNA was isolated using RNeasy micro kit (74104, Qiagen) in combination with QIA shredder columns (79656, Qiagen) according to manufacturer’s instructions including on-column DNAse treatment. RNA concentration and quality was assessed using a Nanodrop 2000 spectrophotometer (Thermo Fisher Scientific). 300 ng of RNA was reverse transcribed using SuperScript IV VILO Master Mix (11756050, Thermo Fisher Scientific) and qPCR was performed in technical triplicates with PowerTrack SYBR Green Master Mix (A46012, Thermo Fisher Scientific). Cq values were normalized using GAPDH as a housekeeping gene.

For standard RT-PCR, 1 µl cDNA was combined with respective primers and OneTaq Quick-Load 2X Master Mix (M0486S, New England Biolabs). For detecting OTX2 splice variants the annealing temperature was 60°C. Primers are summarized in Table 1.

**Table 1:**
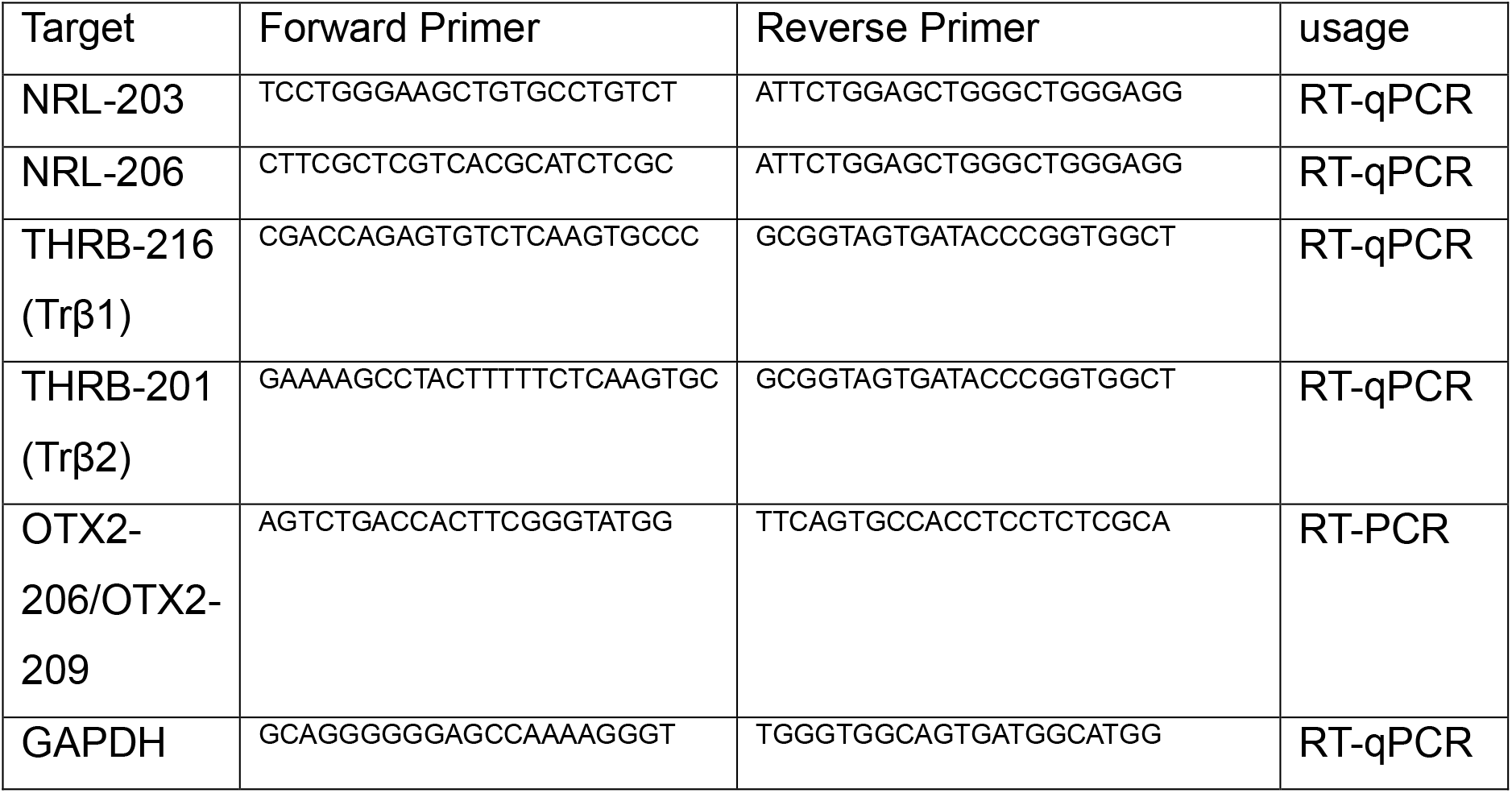
Primers used for PCR.

## Data Records

The dataset contains single cell sequencing of full-length cDNA from human fetal retina of post conception day (PCD) 82. One sample was prepared from central temporal (including the fovea) and one from nasal peripheral retina. Sequencing was performed on an Oxford Nanopore PromethION and deposited at GEO with the accession number GSE342534. It contains the basecalled, unaligned bam files of the samples (M2950_SCR251N_unmapped.bam (nasal.peripheral), M2949_SCR250T_unmapped.bam (central temporal)). Additionally, the corresponding count matrices (mtx format), cell barcodes (tsv format) and feature names (tsv format) for the gene by cell and transcript by cell counts produced by epi2me/wf-single-cell are deposited and are readable by Seurat. The comparison with similar short read data was based on data accessible from GEO under the accession number GSE246169 [16], specifically GSM7864383, GSM7864386, and GSM7864388.

## Data Overview

### LRS sequencing of single cell transcriptomes

Full-length single cell cDNA was generated using 10x Genomics’ Chromium kit according to manufacturer’s protocol and prepared for LRS on an Oxford Nanopore Technologies PromethION. The resulting libraries had a median read length of approximately 700 bp which is slightly lower than the average transcript length of around 1000 bp, but considerably longer than typical 90 bp reads in standard short-read sequencing (Fig. 1A). As expected from polyA capture based RNAseq technologies, we noticed a 3’ bias in the gene body coverage. However, it was less pronounced compared to libraries previously sequenced with short reads [16] as we found more coverage of center and 5’ exons with long reads (Fig. 1B/1C). After filtering and quality control we were left with 6832 cells across 20720 genes and 89919 isoforms. As previously reported [25], most genes (73%) were represented by more than 1 isoform (Fig. 1D). As the gene by cell matrix was generated by independent algorithms (see methods) from the isoform by cell counts, we were interested in how these correlate. We found generally strong Spearman correlation of above 0.5. While the majority of the genes with only 1 isoform showed very strong correlation between isoform and gene counts, genes with multiple detected isoforms had a wider distribution (Fig. 1E).

**Figure 1:**
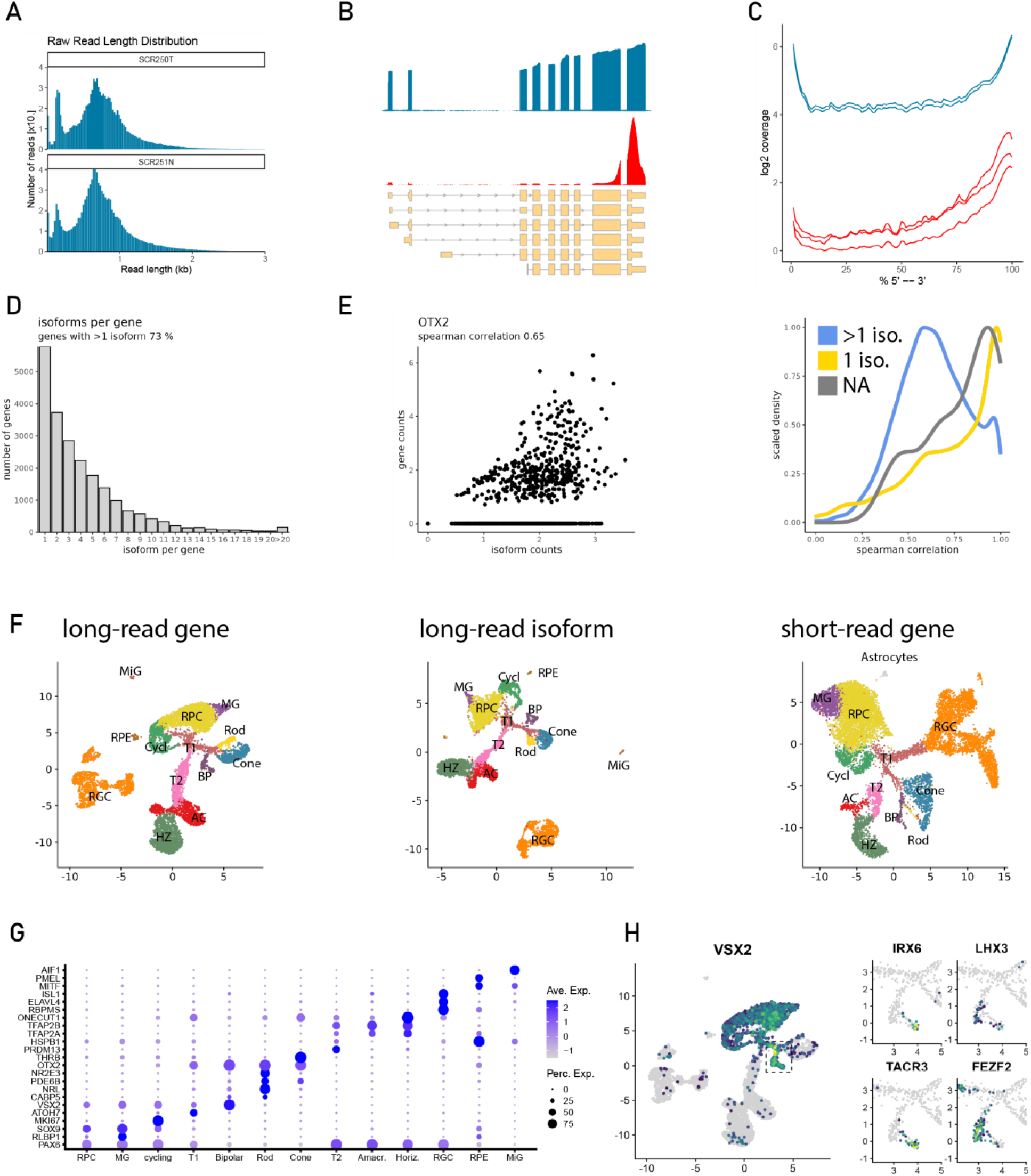
A) LRS produced reads with a median length of approx. 700 bp. B) Gene body coverage across the GAPDH gene and C) across all protein coding genes shows that both LRS and short-read sequencing exhibited a 3’ bias. The bias of LRS was significantly milder as it covered the gene body more evenly. Blue: two LRS samples presented in this study, red: three samples from a published short-read dataset [16]. D) Most detected genes are represented by more than one transcript isoform. E) Correlation between gene counts and corresponding isoform counts is very strong for genes with only one isoform (yellow) and moderate for genes with multiple isoforms (blue) like OTX2. F) We detected all expected cell types in a branching pattern similar to previous published short-read data of retinal development (right) [16]. This was true for both the gene-based (left) and isoform based (center) embeddings. G) Gene expression of canonical markers used for cell type definition. H)The resolution of our data set reveals not only major cell types but also subtypes. The VSX2 positive bipolar cell cluster splits into two sub branches with expression patterns of types 2 and 3a. MiG: microglia; RPE: retinal pigment epithelium; MG: Müller glia; RPC: retinal progenitor cells; BP: bipolar cells; AC: amacrine cells; HZ: horizontal cells; RGC: retinal ganglion cells; T1/T2: transitional cell states; Cyc: cycling progenitor cells.

## Technical Validation

As the human retina matures from the fovea to the periphery [1], it presents an ideal tissue to study development in its whole range. For example, at intermediate stages of development, like PCD 80, the fetal human retina contains mainly earlier generated cell types, like progenitors, cones and RGCs in the periphery, while the center already starts generating later cells, like bipolar cells and Müller glia. For this study we used retina from a PCD 82 fetus and prepared temporal-central, as well as nasal-peripheral retina for subsequent processing.

### Validation of single-cell transcriptome embedding

Initially, we used the gene and isoform matrices separately to perform dimensionality reduction, embedding and cluster definitions. We then used canonical markers to annotate clusters on the gene level (Fig. 1F, Fig. 1G) and transferred them to the same cells in the isoform layer. We found all expected cell types and a similar cluster topography in both UMAP embeddings, with only minor structural differences for the RGCs (Fig. 1F). In general, the resolution of our data was exceptionally good, showing not only all major cell types and branching as in previously published data (Fig. 1F) [2], [16], [26] but even separating bipolar cell subtypes (Fig. 1H). We were able to capture a bifurcation in the bipolar cell fate (VSX2 positive) with differential expression of subtype specific markers FEZF2, LHX3, IRX6, TAC3 (NK3) that are indicative of types 2 and 3a [1], [27] (Fig. 1H).

### Validation of preferential isoform usage in the human retina

After we had established that the gene expression pattern matches what was expected from this type of tissue, we wanted to validate specific isoform expression in several examples of important retinal genes. We first turned to well described developmental transcription factor NRL, which is specifically expressed in rods and essential for their development [1], [28]. This fact is reproduced in our data with highest expression in the rod cluster and some interspersed positive cells in other photoreceptors, RPC and bipolar cells (Fig. 2A). As we explored transcript specific expression levels, we found that the most abundant isoform expressed in rods is NRL-203 (ENST00000397002) (73.5 % of the combined NRL isoform expression) (Fig. 2B, 2C). In fact, the NRL-203 transcript was originally identified as AS321/clone 321B1 in a cDNA library prepared from adult human retina [29]. It is notable that the “default” transcript described by Matched Annotation from NCBI and EMBL-EBI (MANE select [30]) is NRL-206 (ENST00000561028) (Fig. 2B) which accounts for 12.4 % (Fig. 2C). Interestingly, the sequences of NRL-203 and NRL-206 do not differ in translated exons, but rather in their UTRs, which would result in mRNA regulation rather than in protein diversity (Fig. 2D). A striking difference between the transcripts is the transcription start site (TSS). We were able to use this feature to design primers to specifically quantify the distinct transcripts (Fig. 2E). We prepared cDNA from temporal-central and nasal-peripheral regions of fetal human retina and performed qPCR for NRL-203 and NRL-206. NRL-206 was barely detectable, while NRL-203 showed robust expression with expected pattern: increasing with age, especially in the nasal peripheral region. This confirmed the dominant expression of the NRL-203 isoform in fetal human retina.

**Figure 2:**
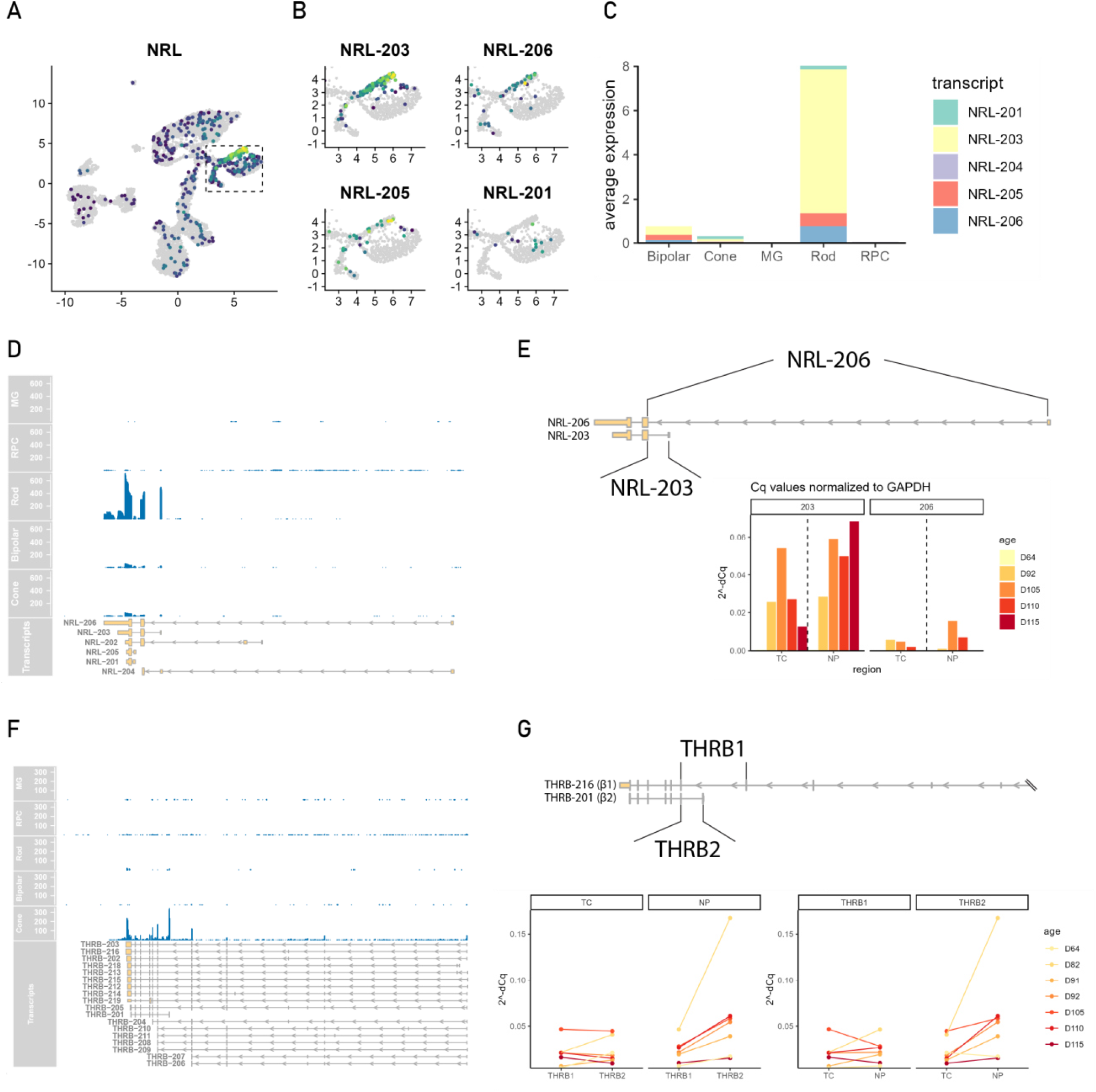
A) NRL gene is primarily expressed in rods with a few positive cells in other clusters. Rods express primarily NRL-203 and to a lower degree NRL-206. C) NRL-203 represents 73 % of total NRL expression in the rod cluster. D) Coverage plot of NRL across different cell types shows rod-specificity and NRL gene structure with a later transcription start site for NRL-203. E) qPCR with primers specific for NRL-203 and NRL-206 confirm the preferential expression of NRL-203 in the temporal center (TC) and nasal periphery (NP) of the human fetal retina. Values normalized to GAPDH expression. Ages are in postconceptional days. F) THRB shows cone specific expression with a preferential coverage of later exons, especially the alternative transcription start site of THRB-201 (Trβ2). G) Isoform-specific primers used for qPCR show that Trβ2 is always higher than Trβ1 in NP and initially in TC, where it switches in samples older than D92 (left). Additionally, the expression differences between regions and thus in developmental time seems to be more dynamic in Trβ2, highlighting the importance to track that isoform specifically (right).

There are genes with different functional isoforms for which a switch is reported during development. It was shown that the Thyroid Hormone Receptor beta (THRB) is important for the development of cones. It has two major protein-isoforms Trβ1 and Trβ2, the latter of which is thought to be specific to only a few regions of the nervous system (including retina). A knockout of Trβ2 in mice led to a switch in opsin expression, from m-opsin to s-opsin, in m-opsin expressing cones [31]. In humans however, the ablation of both isoforms was necessary to achieve a similar result [32]. Furthermore, it was shown that in both mice and humans Trβ2 seems to be dominant in early development (in humans until fetal day 105) while Trβ1 takes over in adult individuals [33]. In humans, the amino acid product of Trβ1 results from multiple annotated transcripts (e.g. ENST00000646209, THRB-216, MANE-select) while Trβ2 is the product of a later TSS yielding fewer exons but a longer peptide (ENST00000280696, THRB-201). Because the base pair differences start to appear only at the 7^th^ exon when counting from the 3’ end, it can be hard to distinguish between the isoforms in short-read sequencing. Our data confirms not only the cone-specific expression of THRB but also the dominant presence of the exon specific for Trβ2 (THRB-201) (Fig. 2F). We performed qPCR on fetal human retina isolated from TC and NP regions from different gestational ages. We found that Trβ2 is always higher than Trβ1 in NP and initially in TC, where it switches in samples older than D92 consistent with developmental findings from whole retina [33]. However, there seems to be little difference in Trβ1 between NP and TC, which suggests that the change in Trβ2 is the more dominant driver and thus more important to track (Fig. 2F).

As mentioned above, OTX2 shows expression that is limited to the photoreceptor and bipolar cell branch (Fig. 3A). Looking at isoform level resolution, we saw that most transcripts are equally distributed among rods, cones and bipolar cells. However, OTX2-206 (ENST00000555006) exhibited cone specificity (Fig. 3B), with little apparent expression in the rod cluster or bipolar cells. Comparison between OTX2-206 and the MANE-select transcript OTX2-209 (ENST00000672264) not only shows the complete absence of 3’ UTR and a different 5’ UTR structure in OTX2-206, but also the exclusion of 8 amino acids (GPWASCPA) in the central translated exon (exon 4 of OTX2-209/exon 3 of OTX2-206) (Fig. 3C). Looking at coverage, it seems that the shorter exon is preferentially expressed. We designed primers spanning the exon junction and the missing amino acids which are predicted to result in bands showing a difference in 24 base pairs. RT-PCR of fetal human retina and RPE confirmed the presence of both alternative exons with a stark bias towards the shorter version. (Fig. 3D). This confirmed similar reports that the shorter OTX2 variant is dominant in mouse retina and RPE [6], [34].

**Figure 3:**
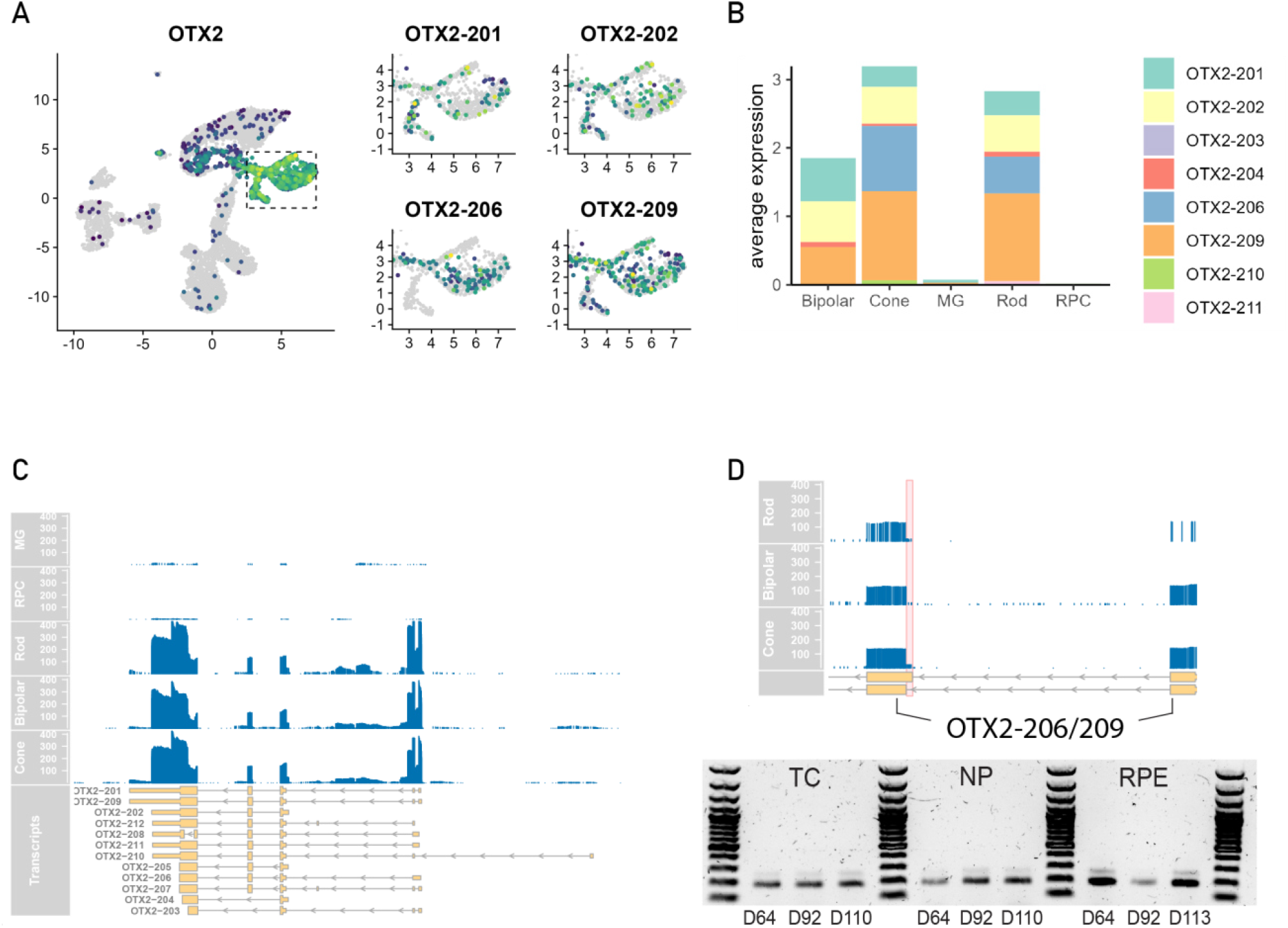
A/B) OTX2 gene is predominantly expressed in photoreceptors and bipolar cells. Most isoforms show distributed expression across these cell clusters, however OTX2-206 seems to be specific for cones. C) Coverage plots show complex transcript isoform structure of the OTX2 gene. D) OTX2-206 has a 24 basepairs indel in its third exon compared to OTX2-209 exon 4 (red highlight). RT-PCR with primers for this exon shows a clear bias towards the shorter version independent of retinal region and in RPE confirming the sequencing result.

### Validation of cell type specific miRNA expression

Most miRNAs arise from the transcription of a host gene and are located in introns and are thus present in the form of pri-miRNA, that is polyadenylated and can be captured by poly(dT) oligonucleotides. Multiple miRNAs can be found inside a single host gene or they can even have their own promoter. Their target specific hairpin structure, however, can be far from the 3’ end [35], [36], [37]. So, while the host transcript can be captured and detected, the actual miRNA sequence can be missed due to 3’ bias in short read sequencing. LRS provides a unique ability to evaluate expression of miRNAs. Therefore, we identified miRNA host genes in our data and localized each to cell types. Some of the most abundant species are shown in figure 4 (Fig. 4A). We were able to validate our results in comparing them with data from previous publications where cell type miRNA expression has been determined from other approaches (e.g. in situ hybridization) [38]. In line with earlier reports, MIR124 was expressed most highly in neuronal cell types as compared with Müller glia, RPC or cycling cells. Our analysis also showed that MIR9 expression was restricted to the Müller glia cluster and identified a MIR7 species to be specifically enriched in cones as well as in immature amacrine and horizontal cells (Fig. 4B).

**Figure 4:**
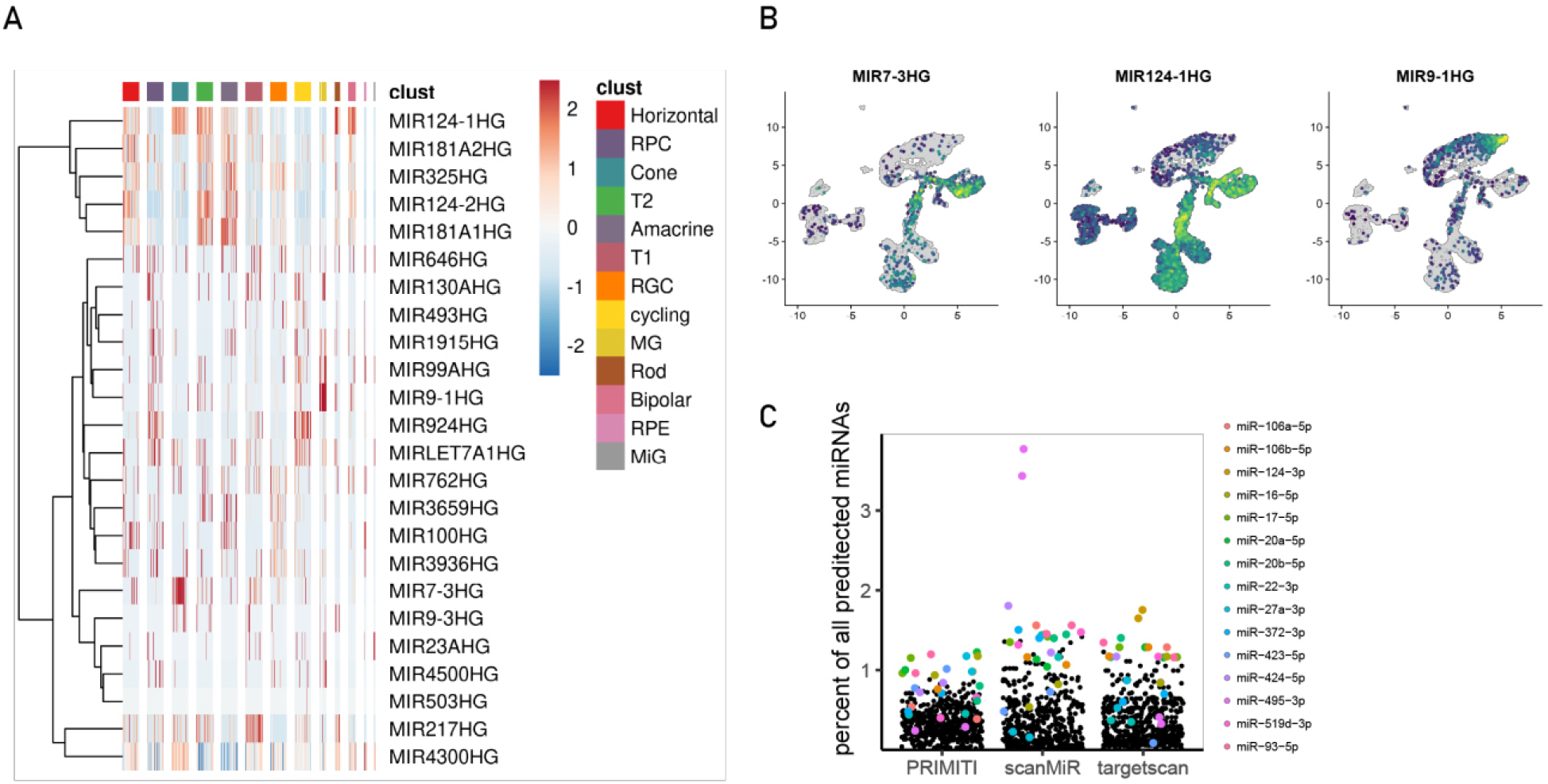
A) Top miRNA host gene expression confirms known retinal miRNA species. B) We found MIR7 to be specifically expressed in photoreceptors, bipolar cells an the transitional state 2, while the well studied MIR124 was specific to all neurons and MIR9 was predominantly expressed in Müller glia. C) Precise knowledge of transcript specific 3’UTRs allowed the prediction of miRNA binding sites. Prediction with three independent tools converged on an overlapping pool of overrepresented miRNAs of which many are known to be important in retinal development. Colored dots represent miRNA species that are in the top 5 overrepresented in any one tool.

### miRNA target site prediction confirms tissue typic regulation

In addition to the information about miRNA expression, the LRS provides information about 3’ UTRs of specific isoforms. As mentioned above, splice isoforms can differ not only in coding regions but to a high degree also in UTRs. The 3’ UTR is especially known to host miRNA binding sites that might lead to transcript degradation or inhibition of translation. We used Targetscan, to identify predictions for conserved miRNA-mRNA pairs [20]. While its predictive power has been appreciated in many studies, the data base has not been updated for several iterations of the GENCODE gene annotation (v19 released in 2013) [20], [39] and exhibits some differences in 3’ UTR definition as compared to the reference (v44 released in 2022) used for our mapping. Therefore, we also used scanMiR which was built on many of the principles of Targetscan and more importantly extended to allow the scanning of custom sequences [22]. Lastly, we used PRIMITI, an orthogonal method which is based on a machine learning model trained with various physicochemical and other features on experimental data [23].

For each miRNA we counted how many transcripts it was predicted to target and related it to the sum of all miRNA-mRNA pairs. So, if any one miRNA accounts for a high percentage of all predicted target pairs, we would consider it overrepresented. For each tool we looked at the top 5 ranked miRNA and saw that these were robustly represented: Any miRNA that was ranked highly in one method was also among the highest ranked in all other methods except for miR-495-3p which was highly ranked in scanMiR and lower in the other. These included miRNAs like miR-124-3p, miR-20a/b-5p or miR-106-a/b-5p that were previously shown to be involved in retinal development [40], [41] (Fig. 4C).

## Data Availability

All raw and processed data was deposited at GEO under the accession number GSE342534. It contains the unmapped bam files as well as the count matrices produced by epi2me-labs/wf-single-cell.

## Code Availability

Code for data processing and figure generation was deposited at https://github.com/rehlab/LR-scRNAseq_fet.hu.retina.

## Acknowledgements

We would like to thank Catherine Ray for excellent technical assistance, Joy Goffena of the lab of Dr. Danny Miller for long-read sequencing related consultation, Dr. Ian Glass from the BDRL for providing human samples and Connor Finkbeiner for productive discussions.

## Author contributions

LK: Conceptualization, Data curation, Formal analysis, Visualization, Writing – original draft, Methodology, Investigation

JP: Investigation

TAR: Funding acquisition, Supervision, Writing – review and editing.

